# Modular mechanisms of immune priming and growth inhibition mediated by plant effector-triggered immunity

**DOI:** 10.1101/2024.02.01.578334

**Authors:** Himanshu Chhillar, Pei-Min Yeh, Hoang Hung Nguyen, Jonathan DG Jones, Pingtao Ding

## Abstract

Excessive activation of effector-triggered immunity (ETI) in plants inhibits plant growth and activates cell death. ETI mediated by intracellular Toll/Interleukin-1 receptor/Resistance protein (TIR) nucleotide-binding leucine-rich-repeat receptors (NLRs) involves two partially redundant signalling nodes in Arabidopsis, EDS1-PAD4-ADR1 and EDS1-SAG101-NRG1. Genetic and transcriptomic analyses show that EDS1-PAD4-ADR1 primarily enhances the immune component abundance and is critical for limiting pathogen growth, whereas EDS1-SAG101-NRG1 mainly activates the hypersensitive cell death response (HR) but is dispensable for immune priming. This study enhances our understanding of the distinct contributions of these two signalling modules to ETI and suggests potential strategies for improving disease resistance in crops without compromising yield.

## Introduction

Inducible plant defence usually involves the concerted action of responses initiated by cell-surface and intracellular immune receptors. Cell-surface pattern recognition receptors (PRRs) initiate pattern-triggered immunity (PTI) upon detection of relatively conserved apoplastic pathogen molecules such as chitin or flagellin. Effector-triggered immunity (ETI) is activated by intracellular nucleotide-binding, leucine-rich repeat receptors (NLRs) upon recognizing specific pathogen effector proteins^1,2^. ETI and PTI together confer a highly effective plant defence against pathogen attack^3–5^. Recent research has significantly expanded our understanding of the signalling pathways downstream of Toll/Interleukin-1 receptor/Resistance protein (TIR)-type NLR (TNL)-triggered ETI, with key roles played by lipase-like proteins ENHANCED DISEASE SUSCEPTIBILITY 1 (EDS1), PHYTOALEXIN DEFICIENT 4 (PAD4), and SENESCENCE-ASSOCIATED GENE 101 (SAG101), and helper NLRs ADR1 (ACTIVATED DISEASE RESISTANCE 1) family and NRG1 (N REQUIREMENT GENE 1) family proteins, which share RESISTANCE TO POWDERY MILDEW 8 (RPW8)-like coiled-coil N-terminal signalling domains^2,6^.

EDS1, PAD4, and SAG101 have emerged as central TNL signalling regulators in *Arabidopsis thaliana* (Arabidopsis hereafter) and many other plant species^7^. These proteins form distinct heterodimeric complexes regulating cell death and disease resistance^8^. EDS1-PAD4 complexes have been shown to play a crucial role in local and systemic acquired resistance (SAR)^9^. Meanwhile, EDS1-SAG101 complexes are hypothesised to be more critical for local immune responses, such as constraining the spread of plant viruses^10^. PAD4 and SAG101 also influence salicylic acid (SA) accumulation, connecting hormonal signalling pathways with the regulation of cell death and disease resistance^11^.

The ADR1 family proteins (ADR1s) in Arabidopsis include ADR1, ADR1-LIKE 1 (ADR-L1), and ADR1-L2, which are also known as helper NLR proteins (hNLRs). They are pivotal in controlling ETI^12^. These proteins act as signal amplifiers, promoting the activation of downstream immune responses upon effector recognition^2,13^. The NRG1 hNLR family proteins (NRG1s), including NRG1A (or NRG1.1) and NRG1B (or NRG1.2), are required by TIR-NLRs to drive hypersensitive response (HR) cell death and contributing to enhanced disease resistance against oomycete pathogens^14^. More recently, it has been shown that *pad4* mutant mimics *adr1s,* while *sag101* mutant mimics *nrg1s* in classical immune responses^15–17^. These genetic results align with the model proposed by several groups that EDS1-PAD4 functions together with ADR1s and contributes more to restricting bacterial growth in TNL-mediated ETI. In contrast, EDS1-SAG101 functions with NRG1 and contributes more towards TNL-mediated cell death induced by ETI^15–20^. These results indicate two distinct immune regulatory nodes associated with EDS1, leading to different downstream signalling.

The growth-defence trade-off, where plants slow growth in response to pests, is a key principle in plant economics affecting both ecosystems and crop breeding^21^. Many constitutive ETI mutants with auto-active NLR have shown severe growth penalties^22^, which might act via various hormonal signalling while enabling plants to thwart pathogen attacks^23^. Recently, it has been shown that constitutive induction of ETI in the absence of PTI can also effectively lead to significant growth arrest^24^. Additionally, prior exposure of plants to pathogens or pathogen-derived ligands primes the plants to induce a much more robust immune response to subsequent encounters^25^. For instance, multiple PAMPs have already been shown to prime Arabidopsis plants against virulent *Pseudomonas syringae* pv. *tomato* (*Pst*) DC3000 bacterium^26^. ETI-induced priming has recently shown a similar effect^24^. However, the contribution of crucial immune regulators in governing the phenomenon of ETI-mediated growth-defence trade-off and immune priming remains inadequately understood.

In this study, we unravel the modular signalling pathways downstream of TNL-induced ETI mediated by EDS1-PAD4-ADR1s and EDS1-SAG101-NRG1s, with particular emphasis on their roles in cell death, immune priming, and ETI-induced growth arrest. We have also explored differential transcriptional outcomes between EDS1-PAD4 and EDS1-SAG101 nodes. This work advances our understanding of plant immunity and sheds light on potential strategies for improving disease resistance in crops without impinging on growth and yield.

## Results

### Differential roles of EDS1-PAD4-ADR1s and EDS1-SAG101-NRG1s in ETI-induced growth arrest

All previous immune phenotypes reported for lipase-like/hNLRs mutants were observed during ETI activation in the presence of PTI. Therefore, it was unknown to what extent prior results are ETI-specific. Here, we used a β-estradiol (E2)-inducible AvrRps4-expressing (‘SETI’) line to activate TNL-mediated ETI in the absence of PTI via the two paired TIR-NLRs RPS4 and RRS1, and RPS4B and RRS1B^24^. When growing SETI plants on an E2-containing medium, we observed growth arrest, including a reduction in size and fresh weight (Figures 1A to 1C). This growth arrest was suppressed entirely in the *eds1-2* mutant (SETI_*eds1*) and the *pad4 sag101* double mutant (SETI_*ps*), indicating a critical role for EDS1, PAD4, and SAG101 in modulating ETI-induced growth inhibition mediated by TNLs.

**Figure 1.**
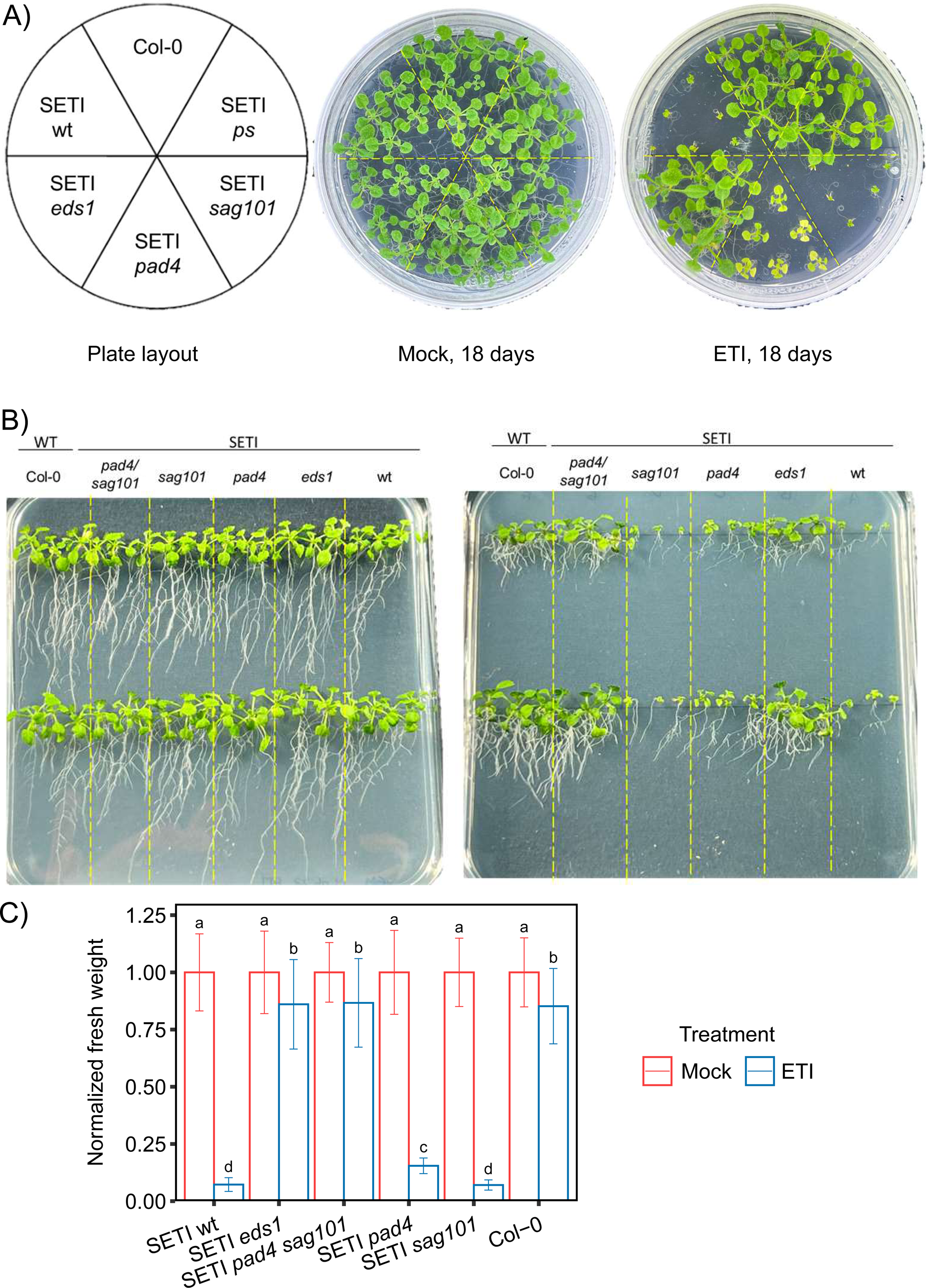
Impact of estradiol treatment on growth arrest phenotype in lipase-like protein mutants. **A**. Indicated mutants of lipase-like proteins (SETI_wt, SETI_*sag101*, SETI_*pad4*, SETI_*pad4sag101*, SETI_*eds1)* were sown on round GM plates with or without E2 (50 μM), and their growth arrest phenotype was recorded 18 days after sowing on E2 plates. (Note: We have used the full name of all the mutants in all the figures for better visualisation) **B**. Indicated mutants of lipase-like proteins were sown on square GM plates with or without E2 (50 μM), and the plates were grown vertically. Phenotypes of growth arrest and lateral root formation were recorded 18 days after sowing on E2 plates. **C.** Indicated mutant lines were sown vertically on square GM plates with or without E2 (50 μM), and the fresh weight was recorded 18 days after sowing on E2 plates. The error bars show the standard deviation. Letters are highlighting statistical differences (LSD-test, p < 0.05)

However, the *sag101* single mutant (SETI_*sag101*) exhibited a growth inhibition pattern like SETI_wt (Figure 1A), implying that SAG101 does not contribute to ETI-induced growth arrest. Interestingly, the *pad4* single mutant (SETI_*pad4*) appeared to partially suppress the ETI-induced growth arrest in the absence of PTI, suggesting PAD4 may play a significant role in mediating ETI-dependent growth arrest independent of SAG101 (Figure 1A). Curiously, a severe inhibition in lateral root formation was observed in SETI_wt and SETI_*sag101,* which was partially relieved in SETI_*pad4* plants and completely relieved in SETI_*eds1* and SETI_*ps* suggesting a role for *PAD4* in controlling root growth and development (Figure 1B).

In parallel, we observed that the *nrg1a nrg1b* double mutant (SETI_*nrg1s*) mirrored the SETI_*sag101* phenotype (Figures S1A to S1C), implying that the NRG1 family proteins may not contribute to growth arrest in response to ETI. Finally, the *adr1 adr1-l1 adr1-l2* triple mutant (SETI_*adr1s*) resembled the SETI_*pad4* phenotype, suggesting a functional correlation between PAD4 and ADR1 proteins in the regulation of ETI-mediated growth arrest.

### Distinct contributions to ETI-mediated cell death by PAD4-ADR1s and SAG101-NRG1s

SETI_wt shows no macro cell death induced by ETI in the absence of PTI^24^. This hypersensitive response (HR) can be measured by recording ion leakage upon E2 induction, which is easily distinguishable at 4 hours post induction (hpi) from the mock treatment and saturates at 22 hpi (Figures 2A, S2E, and S2F). SETI_*eds1* and SETI_*ps* plants do not show any significant increase in ionic conductivity upon E2 treatment, highlighting the interplay of EDS1, PAD4, and SAG101 in mediating the ETI-mediated cell death.

**Figure 2.**
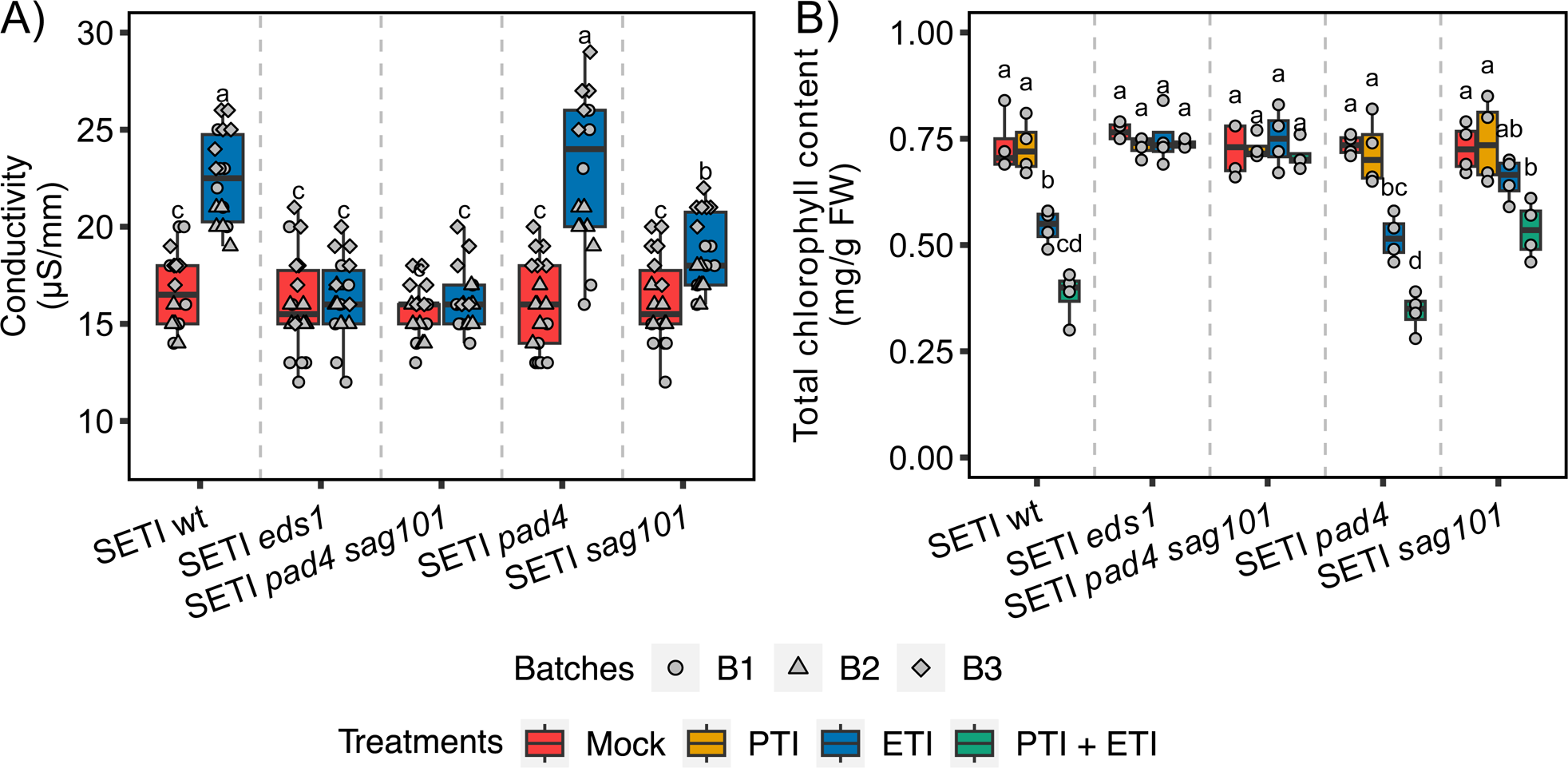
Differential regulation of ETI-induced cell death mediated by lipase-like protein. **A.** Indicated mutants of lipase-like proteins (SETI_wt, SETI_*sag101*, SETI_*pad4*, SETI_*pad4 sag101*, SETI_*eds1*) were infiltrated with E2 or mock and ion leakage was determined at 24 h time point. Letters showing statistical differences (Tukey-HSD-Test, p < 0.05). **B.** Total chlorophyll content was estimated on indicated mutants of lipase-like proteins 3 dpi after E2 infiltration as ETI treatment, *Pst* DC3000 *hrcC*^−^ as PTI treatment, (E2 + *Pst* DC3000 *hrcC*^−^) as PTI+ETI treatment and DMSO dissolved in 10 mM MgCl_2_ as mock. Letters represent statistical differences (Tukey-HSD-Test, p < 0.05).

Interestingly, the increase in ion conductivity in SETI_*pad4* upon E2 induction is similar to SETI_wt, suggesting that PAD4 alone is insufficient to activate ETI-induced cell death (Figure 2A). However, the increase in ion conductivity upon E2 treatment is significantly compromised in SETI_*sag101* plants, showing an independent role of SAG101 from PAD4 in mediating ETI-mediated cell death in the absence of PTI (Figure 2A). Furthermore, SETI_*ps* shows a further reduction in ion leakage as compared to SETI_*sag101*. This suggests a synergistic contribution of PAD4 and SAG101 to cell death (Figure 2A). We have also observed similar trends in the *hNLR* mutants in the SETI background (SETI_*nrg1s* and SETI_*adr1s*) (Figure S2A and S2F). The SETI_*adr1s* mimics SETI_*pad4* while SETI_*nrg1s* mimics SETI_*sag101,* showing that the SAG101-NRG1 node might contribute more to ETI-induced cell death while the PAD4-ADR1 node merely acts as a synergistic module in the presence of the parallel SAG101-NRG1 node.

HR has been shown to have a close association with chlorophyll catabolism^27,28^. Therefore, we sought to test the chlorophyll content as an additional indicator of cell death. A reduction in total chlorophyll content was observed for SETI_wt and SETI_*pad4* upon E2 treatment (Figures 2B and S2C), which was intensified significantly upon the ‘PTI+ETI’ (PTI plus ETI) treatment (50 μM of E2 and *Pst* DC3000 *hrcC*^−^) which is in line with the previous report^4^. However, in SETI_*sag101*, a significant reduction of the total chlorophyll content is observed after ‘PTI+ETI’ treatment but not after PTI nor ETI alone (Figure 2B). This reduction is less significant than in SETI_*pad4* and SETI_wt (Figure 2B). As expected, there is no reduction in chlorophyll content in SETI_*ps* lines. SETI_*adr1s* also showed a significant reduction in chlorophyll content upon ETI and ‘PTI+ETI’ treatments (Figures S2B and S2F), and this reduction is compromised in SETI_*nrg1s*, which is also in agreement with the ion leakage results that the SAG101-NRG1 node contributes more to ETI-induced cell death.

### ETI-mediated immune priming requires both PAD4-ADR1s and SAG101-NRG1s nodes

Pretreatment of SETI_wt plants with E2 shows increased disease resistance against subsequent bacterial infection, indicating ETI alone can prime plant immunity^24^. Consistent with this previous report, we also observed a significant reduction in bacterial growth compared to mock control when SETI_wt plants were pre-infiltrated with E2 one day before being inoculated with the virulent bacterial strain, *Pst* DC3000 carrying an empty vector (Figure 3). In contrast, E2-pretreated SETI_*eds1* and SETI_*ps* plants show complete loss of priming.

**Figure 3.**
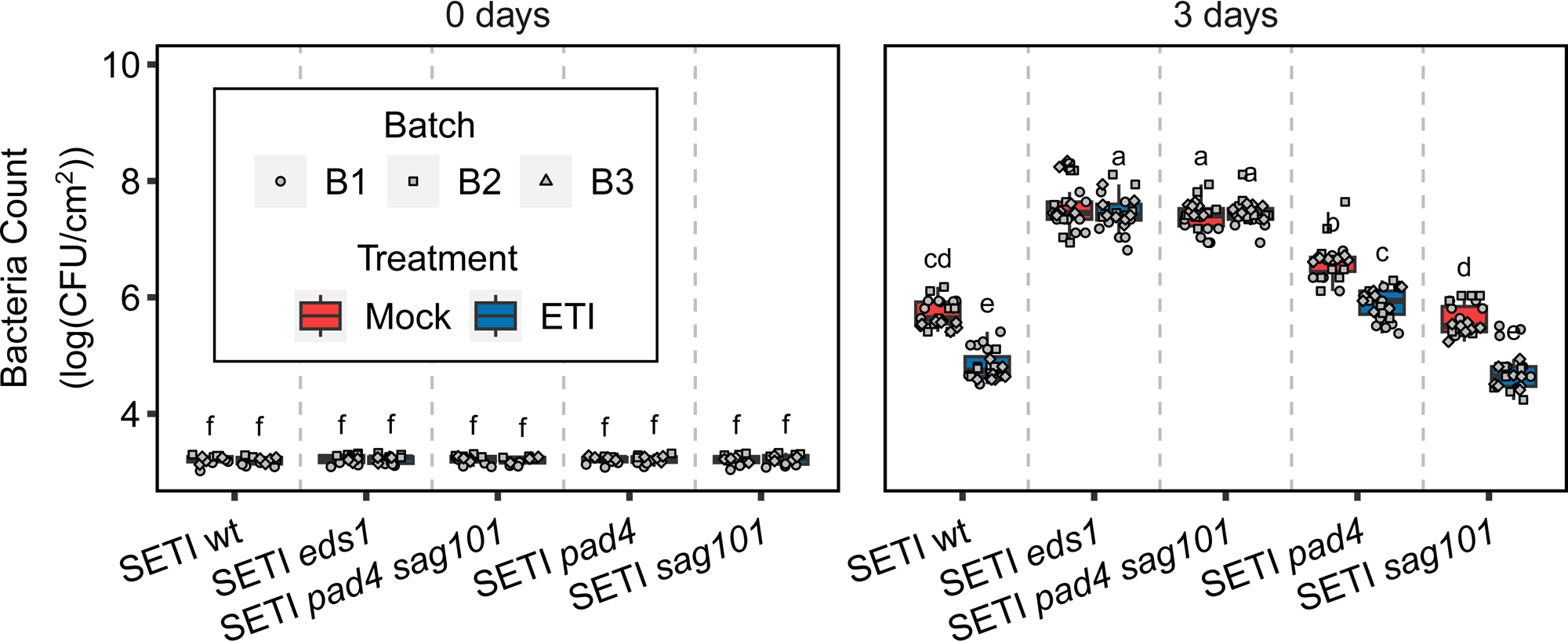
E2-directed disease priming in SETI lipase-like protein mutants. Indicated mutants of lipase-like proteins were infiltrated with E2 or mock one day prior to infiltration with *Pst* DC3000 EV. Bacterial CFU were counted at day ‘0’ and day ‘3’ after infiltration with DC3000 EV. Letters showing statistical differences (Tuckey-HSD-Test, p < 0.05).

In parallel, we then performed an E2- or ETI-induced disease priming assay with SETI_*pad4* and SETI_*sag101* mutant plants. We found that the E2-pretreated SETI_*sag101* plants show no significant difference from SETI_wt plants in disease resistance priming (Figure 3). Similarly, such a priming effect is retained in SETI_*pad4*, though overall bacterial growth in E2-pretreated SETI_*pad4* plants is more than those in SETI_wt and SETI_*sag101* plants (Figure 3). It is noteworthy that the *pad4* but not *sag101* mutant was shown to be partially compromised in PTI-primed disease resistance with the elicitor of a 22-amino-acid epitope from bacterial flagellin and fully compromised in nlp20-induced priming^29^, indicating that PAD4 may play a more significant role in both ETI- and PTI-primed disease resistance than SAG101 in Arabidopsis. However, both PAD4 and SAG101 are required for the disease priming of TNL-mediated ETI.

We further tested ETI-mediated disease priming with SETI_*ngr1s* and SETI_*adr1s* mutants and the higher-order SETI_*helperless* mutant, referring to as a quintuple mutant with the loss-of-function of both *adr1s* and *nrg1s*, *adr1 adr1-l1 adr1-l2 nrg1a nrg1b*^15,17^. E2-pretreated SETI_*nrg1s* show no significant defects in disease priming, which is similar to SETI_wt and SETI_*sag101*, whereas SETI_*adr1s* resembles the phenotype of SETI_*pad4* (Figure S3). A complete loss in disease priming is only observed in E2-pretreated SETI_*helperless* plants, and together with the results from SETI_*ps* (Figures 3 and S3), it can be inferred that both PAD4-ADR1 and SAG101-NRG1 nodes are required for TNL-activated ETI-mediated immune priming.

### ETI-specific defence gene profiling mediated by PAD4-ADR1s and SAG101-NRG1s

We performed genome-wide RNA-seq on SETI mutant lines by specifically inducing ETI through E2 treatment to investigate the differences and similarities of defence gene activation between SAG101-NRG1s and PAD4-ADR1s nodes. We found 5292 differentially expressed genes (DEGs) across all tested genotypes and treatments (Figure 4A, Tables S2-S12). There are 1,902 up-regulated genes in the SETI_wt, 1,707 up-regulated genes in the SETI_*pad4*, and 2,281 up-regulated genes in the SETI_*sag101* E2-treated samples compared to mock-treated samples (Figure S4). The DEGs were substantially reduced in the SETI_*pad4* mutants but not in SETI_*sag101,* indicating that PAD4 plays a major role in TNL-mediated ETI-induced transcriptional reprogramming and its loss-of-function could not be compensated by SAG101. On the other hand, there is no significant reduction in DEGs in SETI_*sag101,* suggesting that PAD4 can largely compensate for the loss-of-function of SAG101. These results highlight the unequal redundancy between PAD4-ADR1s and SAG101-NRG1s, consistent with previous reports that ADR1s can compensate for NRG1s but not *vice versa*^17^. There are no DEGs in SETI_*eds1* and SETI_*pad4 sag101* after E2 treatment compared to mock, demonstrating that *eds1* mutant and *pad4 sag101* double mutant can completely block the TNL-mediated ETI-specific transcriptional reprogramming.

**Figure 4.**
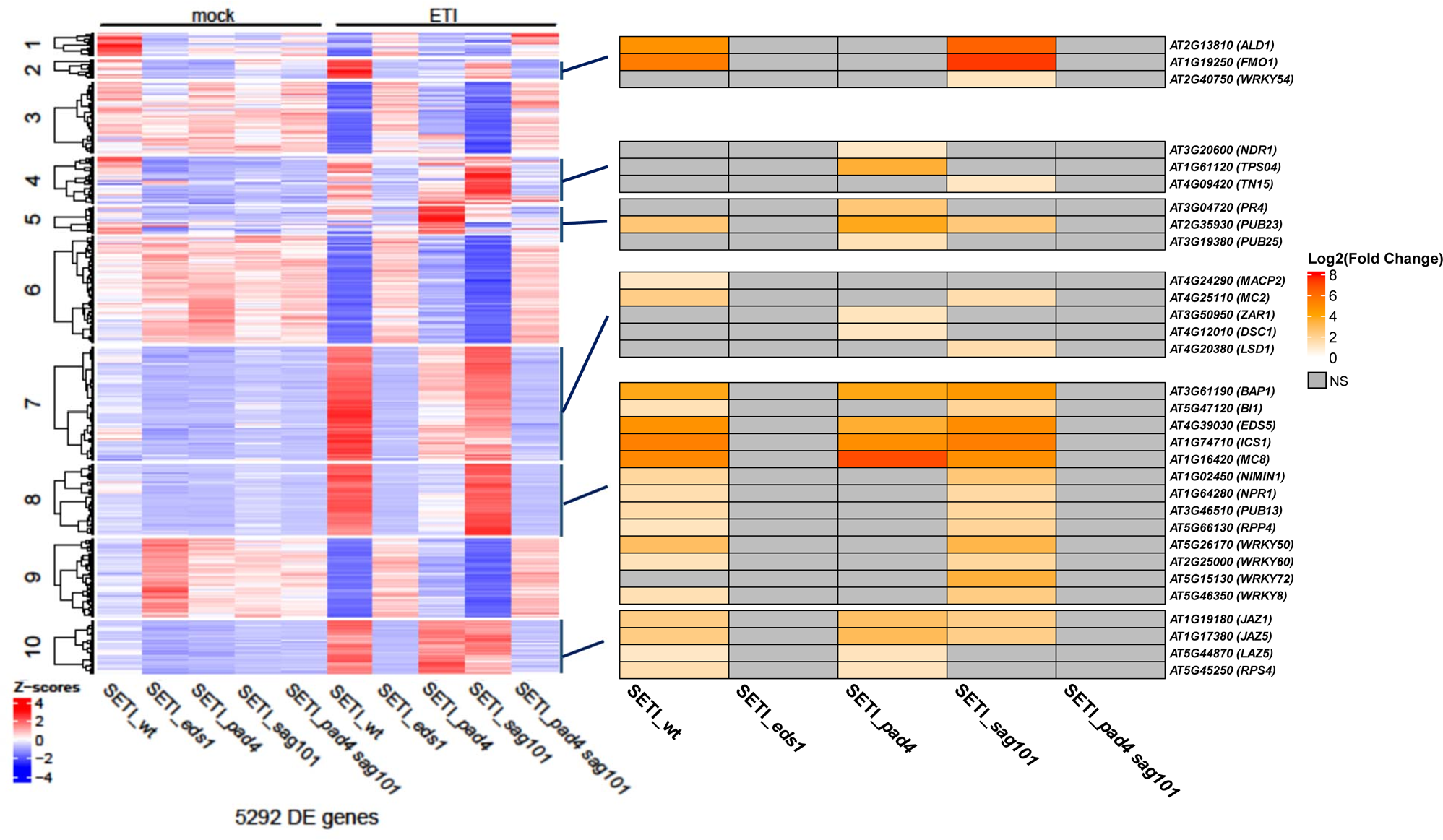
Analysis of TNL-dependent transcriptional changes across SETI lipase-like protein mutants. Five- to six-week-old SETI_wt, SETI_*eds1*, SETI_*pad4*, SETI_*sag101*, and SETI_*pad4 sag101* plants were infiltrated with mock (10 mM MgCl_2_), and 50 μM E2 (ETI). The left heatmap shows the normalized expression z-score of all the differently expressed genes with a false discovery rate (FDR) of p < 0.01 and a fold change of above 1. No significant gene expression changes in SETI_*eds1* and SETI_*ps* after ETI-treatment induction. SETI_wt, SETI_*pad4*, and SETI_*sag101* show visible expression after ETI-treatment induction. The right heat map represents the expression pattern of key immune genes from each immunity-related cluster among the SETI lipase-like protein mutants i.e. SETI_wt, SETI_*pad4*, and SETI_*sag101*.

Hierarchical clustering of the DEGs in the heat map revealed 10 clusters using a Euclidean distance and ward.D clustering algorithm (Figure 4)^30^. Clusters 2, 4, 5, 7, 8, and 10 show defence-related DEGs according to the gene ontology (GO) analysis from g:Profiler (FDR < 0.05, Benjamini-Hochberg) indicating that these are immune-related clusters (Figure 4). The well-known immune-related genes from each cluster and their expression patterns across the SETI mutants were highlighted (Figure 4, Table S1). Among the immune clusters, cluster 2 comprises genes such as the N-hydroxy pipecolic acid (NHP) biosynthetic genes *FMO1* and *ALD1*^31^, while cluster 7 contains genes related to cell death. On the other hand, cluster 8 contains genes mostly related to salicylic acid biosynthesis and signalling, like *ICS1*, *EDS5*, and *NPR1*^32^. Cluster 9 contains genes related to jasmonate signalling.

### Defining the specificity of PAD4- and SAG101-mediated transcriptional reprogramming

The upregulated DEGs were further classified into PAD4- and SAG101-dependent genes and DEGs shared by PAD4 and SAG101 (Table 1). 438 DEGs are PAD4-dependent and up-regulated in both SETI_wt and SETI_*sag101* but downregulated or show no difference in SETI_*pad4*. 85 DEGs are SAG101-dependent and up-regulated in both SETI_wt and SETI_*pad4* but downregulated or show no different in SETI_*sag101* (Figure 5A). Furthermore, 1,258 DEGs show upregulation in SETI_wt, SETI_*pad4*, and SETI_*sag101,* indicating the transcriptional regulation of these DEGs is ETI-specific but redundantly regulated by PAD4 and SAG101 (Figure 5A and S6A to S6C). GO terms related to immune-related biological function were highlighted (Figures 5B to 5D, Tables S12-S14). Interestingly, more DEGs are PAD4-dependent than SAG101-dependent (Figures 5B and 5C). Some genes related to SAR, such as *FMO1* and *ALD1,* depend exclusively on PAD4, while others depend on PAD4 and SAG101 (Figures 5B and 5D, Tables S12 and S14).

**Figure 5.**
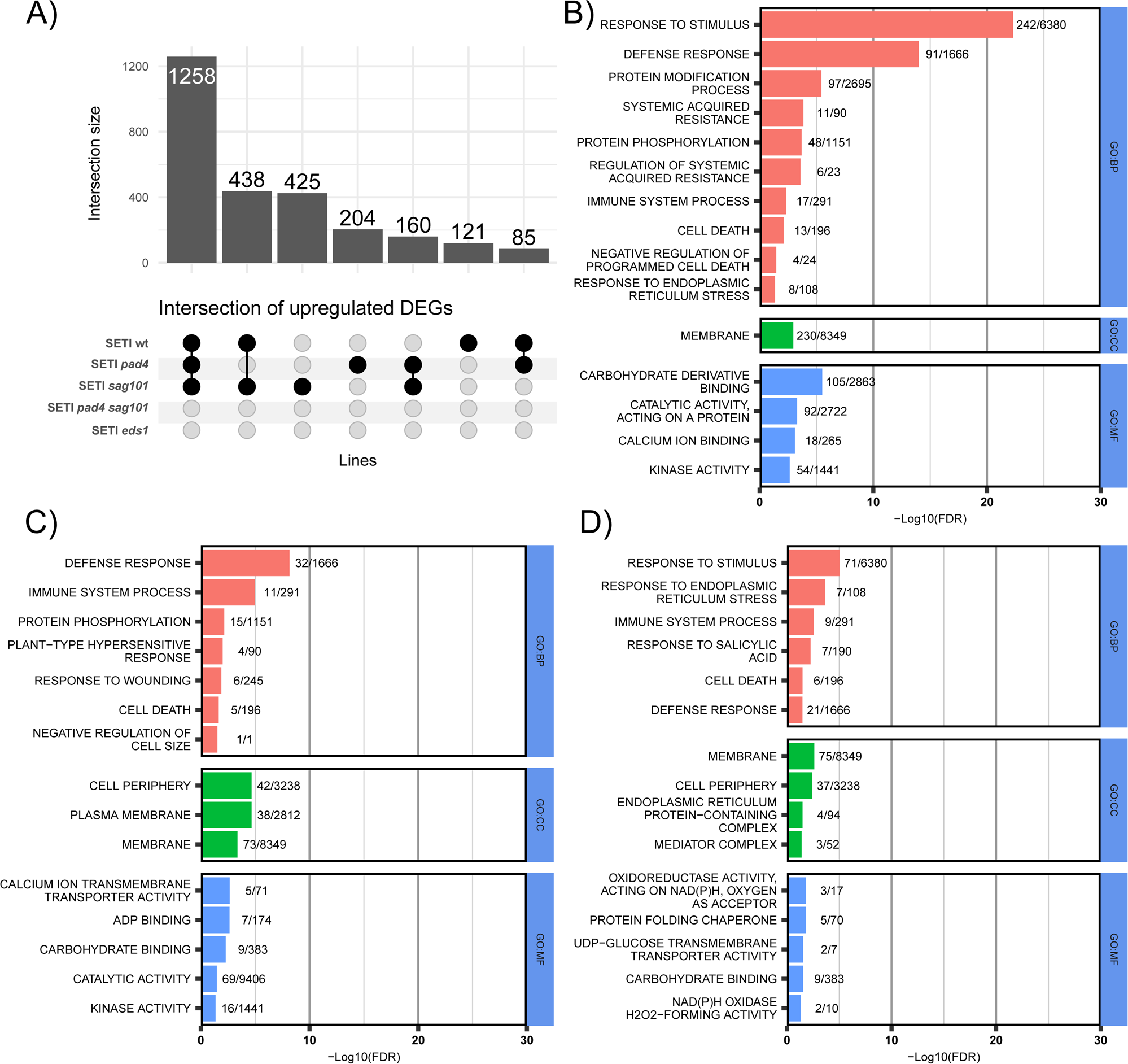
Comparison of gene up-regulation in ETI treatments across lipase-like protein mutants. (A) Up-Set plot shows 1258 DEGs affected by both PAD4 and SAG101. 438 DEGs are *PAD4*-dependent, and 85 DEGs are SAG101-dependent. (B) GO enrichment for PAD4-dependent DEGs. (C) GO enrichment for SAG101-dependent DEGs. (D) GO enrichment for PAD4/SAG101-shared DEGs. GO analysis is done with g:Profiler and the GO terms were shown in different enrichment analyses including biological process (GO:BP), cell components (GO:CC), and molecular function (GO:MF) (FDR = 0.05, Benjamini-Hochberg).

**Table 1.**
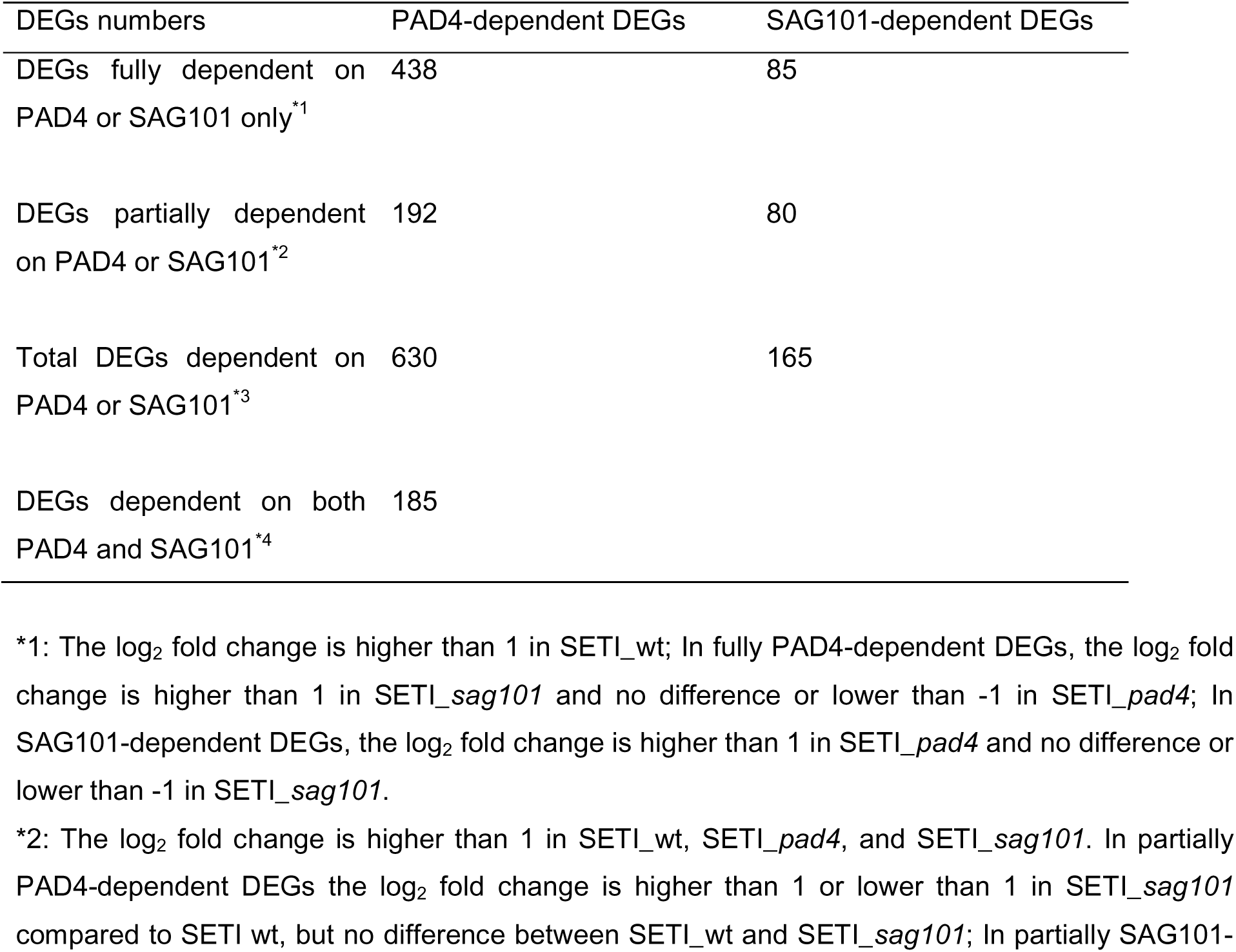
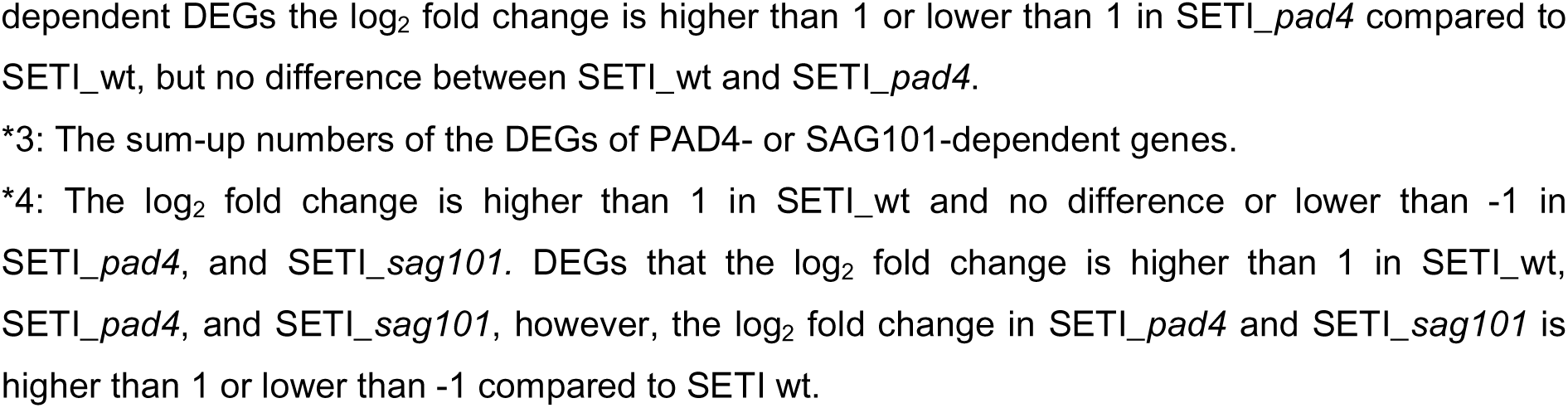
Criteria for sorting the genes as PAD4-dependent, SAG101-dependent, and PAD4/SAG101 dependent.

| DEGs numbers | PAD4-dependent DEGs | SAG101-dependent DEGs |
| --- | --- | --- |
| DEGs fully dependent on PAD4 or SAG101 only<sup>*1</sup> | 438 | 85 |
| DEGs partially dependent on PAD4 or SAG101<sup>*2</sup> | 192 | 80 |
| Total DEGs dependent on PAD4 or SAG101<sup>*3</sup> | 630 | 165 |
| DEGs dependent on both PAD4 and SAG101<sup>*4</sup> | 185 |  |

\*1: The log<sub>2</sub> fold change is higher than 1 in SETI\_wt; In fully PAD4-dependent DEGs, the log<sub>2</sub> fold change is higher than 1 in SETI\_sag101 and no difference or lower than -1 in SETI\_pad4; In SAG101-dependent DEGs, the log<sub>2</sub> fold change is higher than 1 in SETI\_pad4 and no difference or lower than -1 in SETI\_sag101.
\*2: The log<sub>2</sub> fold change is higher than 1 in SETI\_wt, SETI\_pad4, and SETI\_sag101. In partially PAD4-dependent DEGs the log<sub>2</sub> fold change is higher than 1 or lower than 1 in SETI\_sag101 compared to SETI wt, but no difference between SETI\_wt and SETI\_sag101; In partially SAG101-
dependent DEGs the log<sub>2</sub> fold change is higher than 1 or lower than 1 in SETI\_*pad4* compared to SETI\_wt, but no difference between SETI\_wt and SETI\_*pad4*.
\*3: The sum-up numbers of the DEGs of PAD4- or SAG101-dependent genes.
\*4: The log<sub>2</sub> fold change is higher than 1 in SETI\_wt and no difference or lower than -1 in SETI\_*pad4*, and SETI\_*sag101*. DEGs that the log<sub>2</sub> fold change is higher than 1 in SETI\_wt, SETI\_*pad4*, and SETI\_*sag101*, however, the log<sub>2</sub> fold change in SETI\_*pad4* and SETI\_*sag101* is higher than 1 or lower than -1 compared to SETI wt.

It is evident from the ion leakage data that SAG101 contributes more to cell death, but PAD4 has synergistic effects in addition to regulating cell death. As expected, cell-death-related genes were found in all categories, including PAD4-specific, SAG101-specific, and PAD4/SAG101-shared. For instance, genes enriched in cell death-related GO terms such as *METACASPASE 2* (*MC2*), *MC8*, *CYSTEINE-RICH RECEPTOR-LIKE KINASE 13* (*CRK13*), *BAX INHIBITOR 1* (*BI1*) are found to be dependent on PAD4 (Figure 5B, Table S12). On the other hand, genes like *LAZARUS 5* (*LAZ5*) and *CRK4* are dependent on SAG101 (Figure 5C, Table S13), while *BCL-2 ASSOCIATED ATHANOGENE 6* (*BAG6*) and *MEMBRANE ATTACK COMPLEX/PERFORIN-LIKE 2* (*MACP2*) depend on both PAD4 and SAG101 (Figure 5C, Table S14). Although there are DEGs shared by PAD4 and SAG101, many DEGs are regulated solely by either PAD4 or SAG101 (Figure 5A), which sheds light on their functional specificity and unequal redundancy. In summary, PAD4 seems to play a prominent role in regulating ETI-activated transcriptional reprogramming compared to SAG101, which may explain why PAD4 contributes more to the ETI-induced growth arrest and immune priming, whereas SAG101 plays a more prominent role in cell death. It is still unclear how the specificity of transcriptional regulation is achieved by these two unequally redundant nodes downstream of ETI mediated by TNLs.

## Discussion

EDS1 interacts with either of two other lipase-like proteins, PAD4 and SAG101, to initiate the ETI downstream signalling after sensor TNL activation^8,33,34^. Upon TNL activation, helper NLRs ADR1 or NRG1 interact with the EDS1-PAD4 and EDS1-SAG101 modules, respectively^16^. ETI promotes interaction between NRG1 and EDS1/SAG101 but NRG1/EDS1/SAG101 oligomerization requires PTI^19,35,36^. EDS1-PAD4-ADR1s and EDS1-SAG101-NRG1s have been shown to play unequally redundant roles in plant immune responses^16,17^. EDS1/PAD4/ADR1s contribute more to disease resistance, while EDS1/SAG101/NRG1s, contribute more towards cell death. However, the downstream components and the molecular machinery underlying the unequal roles are still not well understood.

Here, we focused on investigating ETI-specific roles of the PAD4 and SAG101 nodes. We use the mutants of EDS1 family proteins and helper NLRs in an inducible ETI genetic background to determine the involvement of these proteins in mediating ETI-specific signalling. A partial relief of growth reduction, as well as of inhibition in the lateral root, was observed in SETI_*pad4* compared to SETI_*sag101* on E2 plates, indicating that PAD4 plays an important role in coordinating the balance between growth, development, and defence^23^. SAG101 also plays a synergistic role with PAD4 in the growth-defence trade-off, as only the SETI_*ps* double mutant shows absolutely no growth inhibition. As the detailed mechanisms of growth-defence trade-off remain unexplored, our report regarding the differential roles of PAD4 and SAG101 in mediating ETI-specific growth inhibition provides additional information to help decipher the underlying molecular mechanisms.

We define immune priming as strengthening plant disease resistance after subjecting plants to a priming treatment. Here, we show that SETI_*pad4* (similarly SETI_*adr1s*) and SETI_*sag101* (similarly SETI_*nrg1s*) both show some immune priming upon ETI activation, though both *pad4* (or *adr1s* triple mutant) and *sag101* (or *nrg1s* double mutant) plants are more disease susceptible^14,16^. These indicate that plants with partially compromised immunity still retain priming capacity for disease resistance. This insight could support improved agricultural practices. For instance, for emerging or remerging pathogens that can partially escape from the immune surveillance system or dampen the immune activation process, one can test whether ETI-mediated immune priming can enhance crop disease resistance. Here, we have only explored the short-term priming capacity in such mutants, and priming could provide longer-term protection. Therefore, it will be interesting to investigate the persistence of this immune memory and how this priming effect has been ‘memorised’ or ‘imprinted’ inside plant cells.

Previous studies have elucidated that HR and disease resistance mechanisms can operate independently^37^. Extending this understanding, our study shows that the SAG101-NRG1s module predominantly facilitates ETI-activated cell death, while the PAD4-ADR1s module is more important for initiating disease resistance. Despite this specificity, both modules have overlapped functions in regulating HR and resistance, though in partially redundant roles. Specifically, NRG1s are required for TNL-mediated disease resistance to obligate biotrophic oomycete pathogens, including the downy mildew agent *Hyaloperonospora arabidopsidis* (*Hpa*) and the white rust causative *Albugo candida*^14^. ADR1s are also required for TNL-mediated resistance against *Hpa*^12^. While the *sag101* single mutant exhibits only isolated single-cell HR cell death, which appears to confer a less severe susceptibility than the *nrg1a nrg1b* double mutant, both SAG101 and NRG1s are consistently implicated as part of the same downstream signalling pathway mediated by EDS1 in multiple instances^14–16,38^. The *pad4* single mutant is capable of producing conidiospores accompanied by single-cell trailing necrosis, which aligns with that observed in the *adr1s* triple mutant^38,39^. More intriguingly, SAG101 and NRG1s appear dispensable for conferring resistance against the hemibiotrophic bacterial pathogen *P. syringae*, where the PAD4-ADR1s node assumes a more important role. Therefore, it is imperative for future studies to comprehensively evaluate disease resistance across a range of mutants, including those carrying mutations in the lipase-like protein-encoding gene (*pad4*, *sag101*, and their double mutant), as well as those affecting hNLR-encoding genes (*nrg1s*, *adr1s*, and their quintuple *helperless* mutant). It is also crucial to expand testing their responses to more different types of plant pathogens. Furthermore, it is key to investigate further how each node modulates the functional preference towards HR and resistance and how these are linked to the differential phenotypes of ETI-induced growth inhibition and disease priming. In addition, it will provide valuable insights into NLR-mediated immune effectiveness to explore how the synergistic effects of SAG101-NRG1s and PAD4-ADR1s nodes are achieved.

Importantly, in real interactions with microbes, ETI is always preceded by PTI, making it difficult to study ETI-specific responses. For instance, in the heat map that encompasses all the treatments (PTI, ETI, ‘PTI+ETI’), gene expression patterns in ‘PTI+ETI’ are much less distinguishable across different lipase-like mutants compared to those in the ETI-only treatment (Figure S5). We suspect that PTI can potentially mask the overall ETI transcriptome, whereas ETI-specific treatment can overcome this. We also found similar effects in several recent reports^17,40^. TNLs or TIR-only proteins possess enzymatic activities to produce small molecules that associate with different EDS1 complexes together with either PAD4-ADR1s or SAG101-NRG1s^41^. Our transcriptome profiling results for either PAD4-ADR1s or SAG101-NRG1s modules specific to ETI might provide additional insights. It has been shown that EDS1-SAG101-NRG1s hNLR resistosome formation requires the cell-surface immune receptor-mediated PTI; it would be interesting to compare the gene expression patterns between PTI, ETI, and ‘PTI+ETI’ in SETI_*pad4*. A few other important unresolved questions remain: How do the resistosomes formed by NRG1s and ADR1s activate ETI downstream responses? Are the cation channel activities of NRG1s and ADR1s sufficient to activate the observed transcriptional reprogramming?

More broadly, it has been reported that PAD4 and SAG101 from different plant species may have various functions during TNL-mediated ETI activation. For instance, the Solanaceae genome mostly encodes two SAG101 isoforms (SAG101a, SAG101b), and EDS1-SAG101b has been shown to play a crucial and sufficient role in nearly all TNL-mediated ETI immune responses in *Nicotiana benthamiana* (*Nb*), while *Nb*PAD4 did not show any significant roles^34^. However, similar to *Nb*SAG101s, *Nb*NRG1 is important in regulating ETI downstream of EDS1 in *Nb*^35,42^. Conceivably, the EDS1-SAG101-NRG1 node plays a crucial role in regulating both disease resistance and cell death in *Nb*, while in contrast, EDS1-SAG101-NRG1 in Arabidopsis is more specialised to HR. It will be interesting in the future to generate ETI-inducible lines in *Nb* that are similar to SETI in Arabidopsis, to study ETI-specific responses mediated by *Nb*PAD4 and *Nb*SAG101 to dissect the ETI-specific downstream signalling in comparison to those in Arabidopsis, which may provide more insights in ETI-mediated growth-defence trade-off and immune priming. In summary, our studies have provided useful datasets to dissect modular mechanisms of immune priming and growth inhibition mediated by ETI, which could underpin smarter plant breeding for disease resistance.

## Materials and methods

### Plant material and growth conditions

*Arabidopsis thaliana* accessions Col-0 and a β-estradiol (E2) inducible Super-ETI line (SETI_wt), were used as the wild-type controls in this study. The lipase-like mutants SETI *eds1-2* (SETI_*eds1*), SETI *pad4-1* (SETI_*pad4*), SETI *sag101-1* (SETI_*sag101*), and SETI *pad4-1 sag101-1* (SETI_*ps*) and helper NLR mutants, including SETI *ad1-1 adr1-L1 adr1-L2 nrg1a nrg1b* (SETI_*helplerless*), SETI *nrg1a nrg1b* (SETI_*nrg1s*), and SETI *adr1-1 adr1-L1 adr1-L2* (SETI_*adr1s*) are generated by crossing all the previously reported mutants with SETI plants^12,14,38^.

For square and round plate assay, seeds were processed by liquid sterilisation (70% ethanol 5 minutes, bleach solution 5 minutes, 100% ethanol 5 minutes, washed three times with sterilised water, and soaked into 0.5% agarose in 4 °C dark condition for one day). Seeds were sown on a 9 cm petri dish or square petri dish (120 mm x 120 mm), GM (Germination media) plate, or GM plate with 50 µM of E2. Plants were grown at 21 °C under long-day conditions (16 h light, 8 h dark), and at 50% humidity. Photos were taken 14 and 18 days after sowing. Col-0 was used as the negative and SETI as the positive control.

### Fresh weight measurement

The seeds were processed by liquid sterilisation and sown on a square petri dish, GM plate, or GM plate with 50 µM of E2 for 18 days. Six seedlings were pooled together to measure the fresh weight. All statistics and figures are generated in R version 4.3.1. ANOVA (p < 0.05) was used for identifying significant factors. A least significant difference (LSD) test (p < 0.05) was used to identify differences between treatment and lines. A detailed statistical summary is available on GitHub: https://github.com/dinglab-plants/ETI_Project.

### RNA-seq raw data processing, alignment, quantification of expression, and data visualisation

DMSO in 10 mM MgCl_2_, 50 μM E2 in 10 mM MgCl_2_, *Pseudomonas fluorescens* (Pf0-1) in 10 mM MgCl_2,_ or Pf0 and 50 μM E2 in 10 mM MgCl_2_ was infiltrated in 5 to 6-week-old Arabidopsis leaves with 1 ml needleless syringe for RNA-seq sample collection. Two leaves per sample were collected at 0 and 4 hpi. RNA was extracted with the Zymo RNA extraction kit ^43^.

The RNA sample was sequenced by Novogene. Raw reads were trimmed into 390 bp clean reads by the Novogene bioinformatics service. At least 12 million single-end clean reads for each sample were provided by Novogene for RNA-seq analysis. All reads passed FastQC before the following analyses^44^. All clean reads were mapped either to the TAIR10 Arabidopsis genome/transcriptome via TopHat2 or to a comprehensive Reference Transcript Dataset for Arabidopsis Quantification of Alternatively Spliced Isoforms (AtRTD2_QUASI) containing 82,190 non-redundant transcripts from 34,212 genes via Galaxy and Salmon tools ^45,46^. The estimated gene transcript counts were used for differential gene expression analysis and statistical analysis with the 3D RNA-seq software ^47^. The low-expressed transcripts were filtered if they did not meet the criteria of ≥3 samples with ≥1 count per million reads. The batch effects between biological replicates were removed to reduce artificial variance with the RUVSeq method ^48^. The expression data were normalised across samples with the TMM (weighted trimmed mean of M-values) ^49^. The significance of expression changes in the contrasting groups ‘SETI_wt_ETI vs SETI_wt_mock’, groups ‘SETI_*eds1*_ETI vs SETI_wt_mock’, ‘SETI_*pad4*_ETI vs SETI_wt_mock’, ‘SETI_*sag101*_ETI vs SETI_wt_mock’ and, ‘SETI_*ps*_ETI vs SETI_wt_mock’ were determined by the limma-voom method ^50,51^. A gene was defined as a significant differentially expressed gene (DEG) if it had a Benjamini–Hochberg adjusted P-value < 0.01 and log2[fold change (FC)] ≥ 1. The GO term analysis was analysed with g:Profiler^52^ (Figure S7).

### Electrolyte leakage assay

Two leaves of 5-week-old Arabidopsis plants were hand infiltrated using a 1-ml needleless syringe with 50 μM E2 dissolved in Mili-Q water or DMSO in Mili-Q water as mock. Leaf discs were collected with a 7-mm diameter cork borer from infiltrated leaves on paper towels. Leaf discs were dried and transferred into 2 ml of deionised water in 12-well plates (2 leaf disks per well. The plate was incubated for 30 minutes in a growth chamber with controlled conditions at 21 °C under long-day conditions (16-h light/8-h dark) with a light intensity of 120-150 µmol m^−2^. The water was replaced after incubation with 2 ml of deionised water. Electrolyte leakage was measured with Pocket Water Quality Meters (LAQUAtwin-EC-33; Horiba) calibrated at 1.41 mS/cm. Around 100 μl of the sample was used to measure conductivity at the indicated time points. ANOVA (p < 0.05) was used for identifying significant factors. Tukey-HSD-Test (p < 0.05) was used to identify differences between treatment and lines. A detailed statistical summary is available on GitHub: https://github.com/dinglab-plants/ETI_Project.

### Bacterial growth assay

*Pst* DC3000 EV (carrying empty vector) was grown on selective King’s B (KB) medium plates containing 15% (w/v) agar, 25 µg ml^−1^ rifampicin, and 50 µg ml^−1^ kanamycin for 48 h at 28 °C. Bacteria were harvested and resuspended in 10 mM MgCl_2_. The suspension concentration was adjusted to an optical density of 0.001 at 600 nm [OD_600_=0.001, representing ∼5×10^5^ colony-forming units (CFU) ml^−1^]. Two 5-week-old Arabidopsis leaves were hand infiltrated with 50 μM E2 dissolved in 10 mM MgCl_2_ or DMSO in 10 mM MgCl_2_ with a needleless syringe. The following day, the same leaves were infiltrated with bacteria. For quantification, leaf samples were harvested with a 7-mm diameter cork borer, resulting in leaf discs with an area of 0.38 cm^2^. Two leaf discs per leaf were collected as a single sample. For each genotype and condition, four samples were collected immediately after infiltration as ‘day 0’ samples, and eight samples were collected at 3 dpi as ‘day 3’ samples to compare the bacterial titres between different genotypes, conditions, and treatments. For ‘day 0’, samples were ground in 200 μl of 10 mM MgCl_2_ and spotted (10 μl per spot) on selective KB medium agar plates to grow for 48 h at 28 °C. For ‘day 3’, samples were ground in 200 μl of infiltration buffer, serially diluted (5, 50, 5×10^2^, 5×10^3^, 5×10^4^, 5×10^5^ times), and spotted (6 μl per spot) on selective King’s B medium agar plates to grow for 48 h at 28 °C. The number of colonies (CFU per drop) was counted, and bacterial growth was represented as CFU cm^−2^ of leaf tissue. ANOVA (p < 0.05) was used for identifying significant factors. Tukey-HSD-Test (p < 0.05) was used to identify differences between treatment and lines. A detailed statistical summary is available on GitHub: https://github.com/dinglab-plants/ETI_Project.

### Chlorophyll content estimation

Two leaves of 5-week-old Arabidopsis plants (SETI_wt, SETI_*eds1*, SETI_*pad4*, SETI_*sag101*, SETI_*ps*, SETI_*adr1s*, SETI_*nrg1s*, and SETI_*helperless*) were hand infiltrated using a 1 ml needleless syringe with 50 μM E2 dissolved in 10 mM MgCl_2_ as ETI treatment, *Pst* DC3000 *hrcC*^−^ (0.2 OD) dissolved in 10 mM MgCl_2_ as PTI treatment, E2+*Pst* DC3000 *hrcC*^−^ with same concentration as ‘PTI+ETI’ treatment and DMSO dissolved in 10 mM MgCl_2_ as mock. Leaf disks were collected using a 7-mm diameter cork borer from 3 individual plants as 1 sample for each treatment. Samples were ground and resuspended in 1 ml of 80% acetone. The samples were then centrifuged at 10,000 g for 5 mins. The absorbance of the supernatant was measured at 645 nm and 633cnm using UV–VIS spectrophotometer (UV-6300PC, VWR). Total and chlorophyll a, b content were calculated according to the following equations ^53^:

> Chl-a = 12.72A663-2.59A645/1000 mg per g FW (mg g^−1^)

> Chl-b = 22.9A645 −4.67A663/1000 mg per g FW (mg g^−1^)

> Chl-t= 20.31 A645 +8.05 A663/1000 mg per g FW (mg g^−1^)

ANOVA (p < 0.05) was used for identifying significant factors. Tukey-HSD-Test (p < 0.05) was used to identify differences between treatment and lines. A detailed statistical summary is available on GitHub: https://github.com/dinglab-plants/ETI_Project.

## Data analysis tools

R version 4.3.1 was used for data analysis. Boxplots and bar plots were generated using ggplot2. The heatmap on the right side in Figure 4 was generated with the R package ‘ComplexHeatmap’^54^. The UpSet and volcano plots were generated with the R packages ‘ComplexUpset’^55^ and ‘EnhancedVolcano’^56^. The following packages were used for data formatting and statistics: ‘Tidyverse’^57^, ‘multcompView’^58^, ‘agricolae’^59^, ‘ggpubr’^60^, ‘stringr’^61^, ‘r2r’^62^ and ‘circlize’^63^. RNA-seq analysis was done as described above.

## Data and code availability

The RNA-seq data for this study have been deposited in the European Nucleotide Archive (ENA) at EMBL-EBI under accession number PRJEB62154. All codes are available via GitHub: https://github.com/dinglab-plants/ETI_Project.

## Author contributions

PD conceptualised and oversaw the inception of the research project. The experimental work was collaboratively conducted by PD, HC, and PY. Data analysis and figure generation were performed by HC, PY, HHN, and PD. JDGJ was involved throughout the entire project, providing valuable discussions that significantly shaped the research. The initial manuscript draft was written by HC and PD. All co-authors contributed to subsequent revisions and editorial processes. The final manuscript was prepared by HC and PD and was approved for submission by all authors.

## Acknowledgements

We thank Marc Gebauer for their technical assistance with the genotyping of mutants and Ewout van Diepen for their invaluable help during the more labour-intensive aspects of our experimentation. Their contributions were instrumental to the successful completion of this work. HC, PY, HHN, and PD acknowledge a European Research Council Starting Grant ‘R-ELEVATION’ (grant agreement: 101039824). JDGJ was supported by the Gatsby Foundation (United Kingdom). We sincerely thank the readers of our initial preprint for their valuable feedback and for pointing out errors. We have incorporated these corrections into this second version. We welcome any further suggestions or comments that can help us improve our manuscript.

---

**Figure S1.**
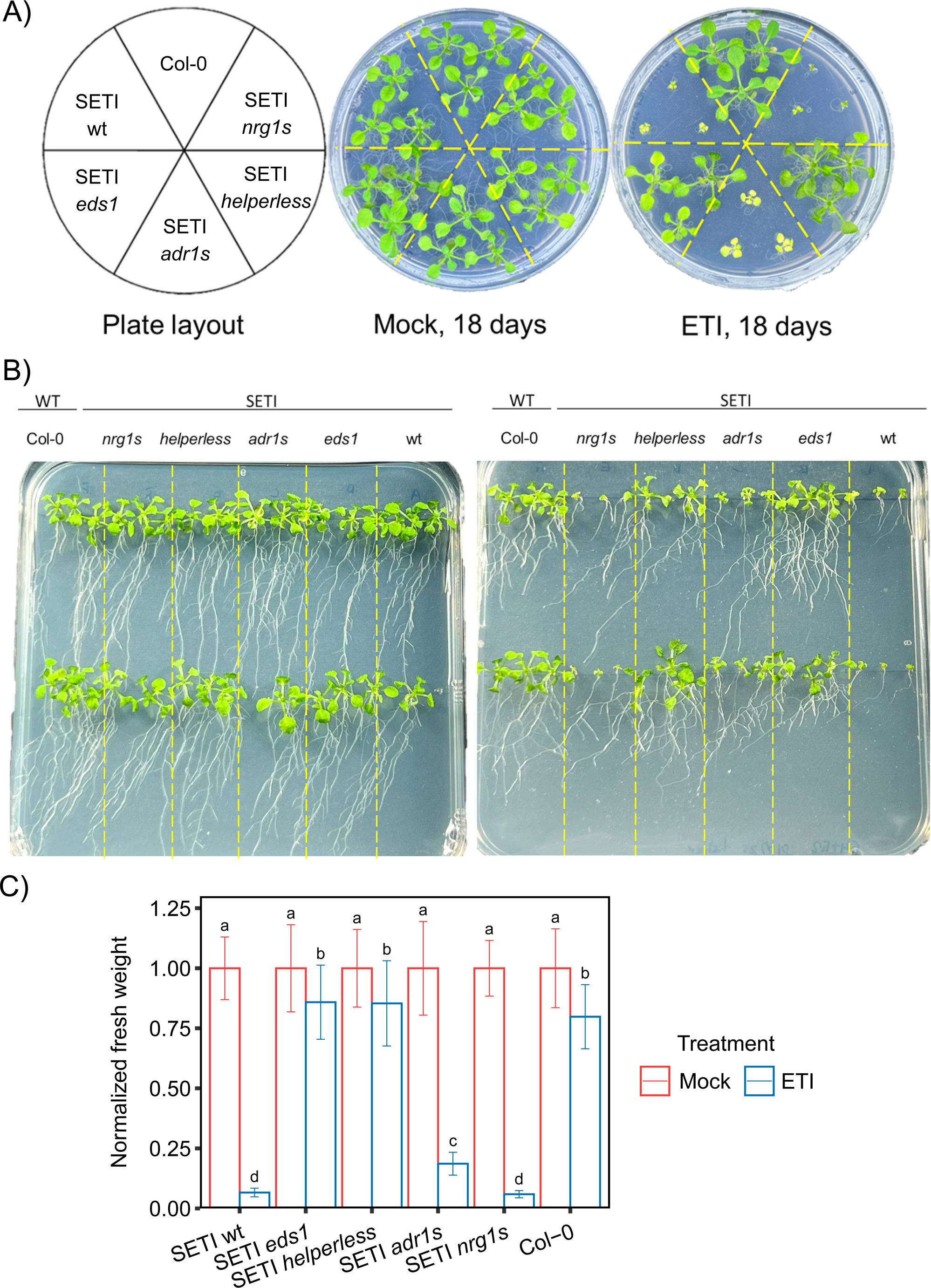
Impact of estradiol treatment on growth arrest phenotype in helper NLR mutants. **A**. Indicated mutants of helper-NLRs (SETI_wt, SETI_*nrg1s*, SETI_*adr1s* and SETI_*helperless,* SETI_*eds1*) were sown on round plates with or without E2 (50 μM), and their growth arrest phenotypes were observed 18 days after sowing on E2 plates. **B.** Indicated helper-NLR mutants were sown on square plates with or without E2 (50 μM) and were grown vertically. Phenotypes of growth arrest and lateral root formation were recorded 18 days after sowing on E2 plates. **C.** Indicated mutant lines were grown on vertical GM plates with or without E2 (50 μM), and the fresh weight was recorded 18 days after sowing on E2 plates. The error bars indicate the standard deviation. Letters showing statistical differences (LSD-test, p < 0.05)

**Figure S2.**
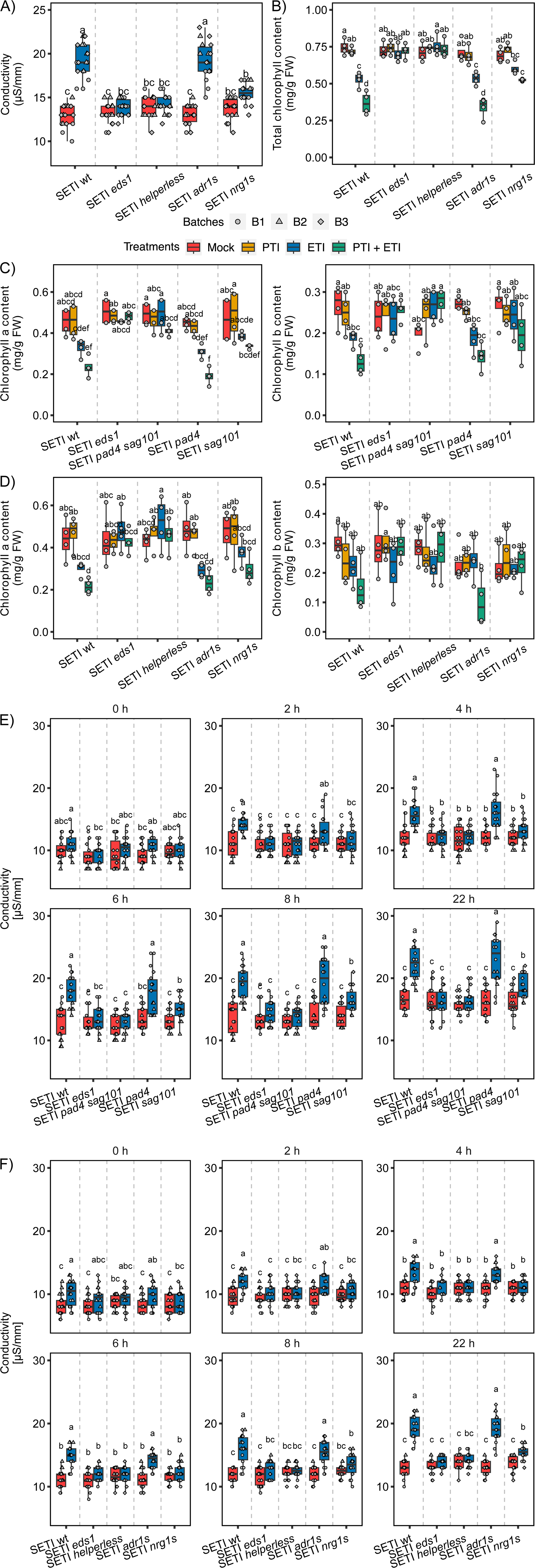
Varying responses SAG101-NRG1s and PAD4-ADR1s modules in ETI-induced cell death. **A.** Indicated mutant lines of helper-NLRs (SETI_wt, SETI_*nrg1s*, SETI_*adr1s*, SETI_*helperless*, SETI_*eds1*) in SETI background were infiltrated with E2 or mock and ion leakage was determined at 24 h time point. Letters showing statistical differences (Tukey-HSD-Test, p < 0.05). **B**. Total chlorophyll content was estimated on indicated mutants of helper NLRs 3 dpi after E2 infiltration as ETI treatment, *Pst* DC3000 *hrcC*^−^ as PTI treatment, (E2 + *Pst* DC3000 *hrcC*^−^) as PTI+ETI treatment and DMSO dissolved in 10 mM MgCl_2_ as mock. Letters showing statistical differences (Tukey-HSD-Test, p < 0.05). **C**. SETI lipase-like protein mutants were hand infiltrated with mock, E2 (ETI), E2 + *Pst* DC3000 *hrcC*^−^ (PTI+ETI) and chlorophyll a and b content was recorded 3 dpi. Letters showing statistical differences (Tukey-HSD-Test, p < 0.05) **D**. SETI helper-NLR mutants were hand infiltrated with mock, E2 (ETI), E2 + *Pst* DC3000 *hrcC*^−^ (PTI+ETI) and chlorophyll a and b content was recorded 3 dpi. Letters showing statistical differences (Tukey-HSD-Test, p < 0.05) **E.** Indicated mutants of lipase-like proteins (SETI_wt, SETI_*sag101*, SETI_*pad4*, SETI_*pad4 sag101*, SETI_*eds1*) were infiltrated with E2 or mock and ion leakage was determined at several timepoints including 0, 2, 4, 6, 8 and 22h. Letters showing statistical differences (Tukey-HSD-Test, p < 0.05). **F.** Indicated mutant lines of helper NLRs (SETI_wt, SETI_*nrg1s*, SETI_*adr1s*, SETI_*helperless*, SETI_*eds1-2*) in SETI background were infiltrated with E2 or mock and ion leakage was determined at several timepoints including 0, 2, 4, 6, 8 and 22h. Letters showing statistical differences (Tukey-HSD-Test, p < 0.05).

**Figure S3.**
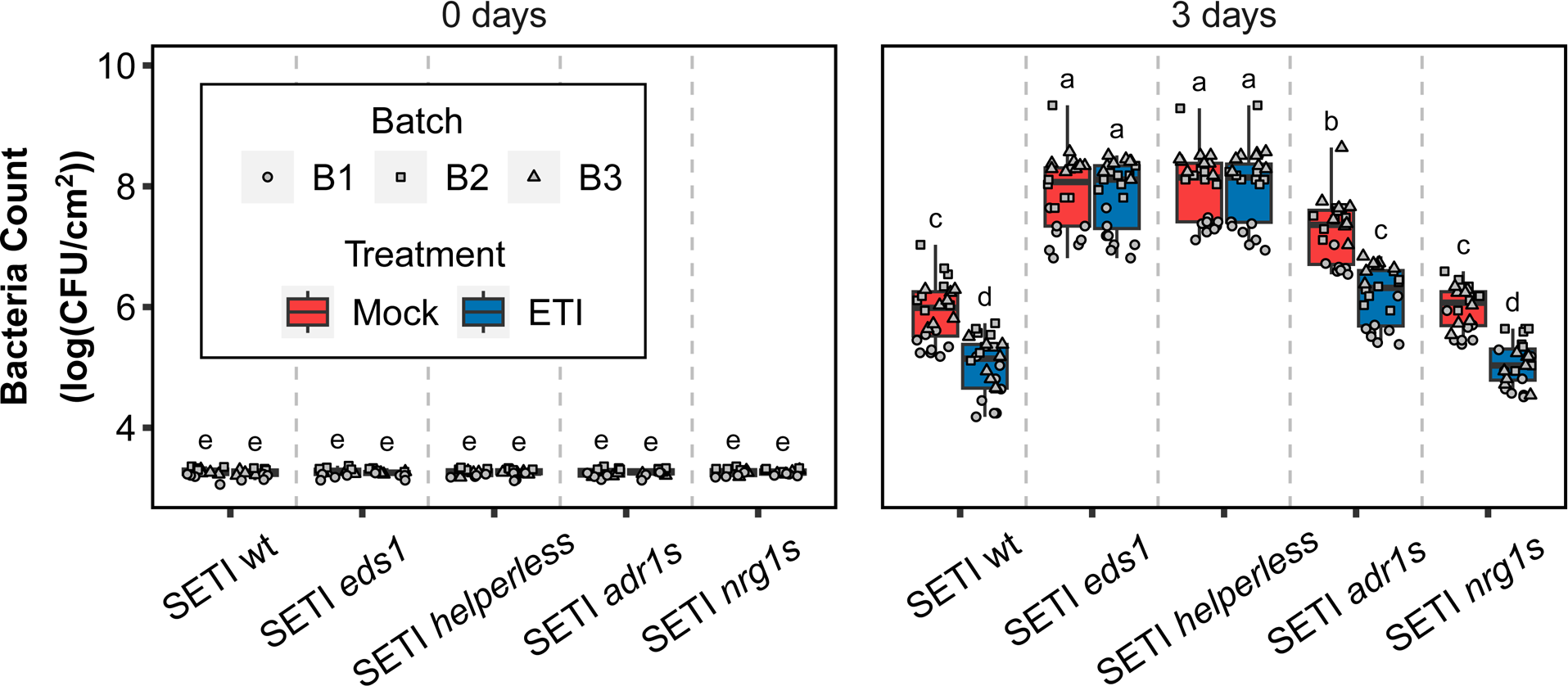
E2-directed disease priming in SETI helper-NLR mutants. Indicated mutants of helper NLRs were infiltrated with E2 or mock one day prior to infiltration with *Pst* DC3000 EV. Bacterial CFU were counted at day ‘0’ and day ‘3’ after infiltration with DC3000 EV. Letters showing statistical differences (Tuckey-HSD-Test, p < 0.05).

**Figure S4.**
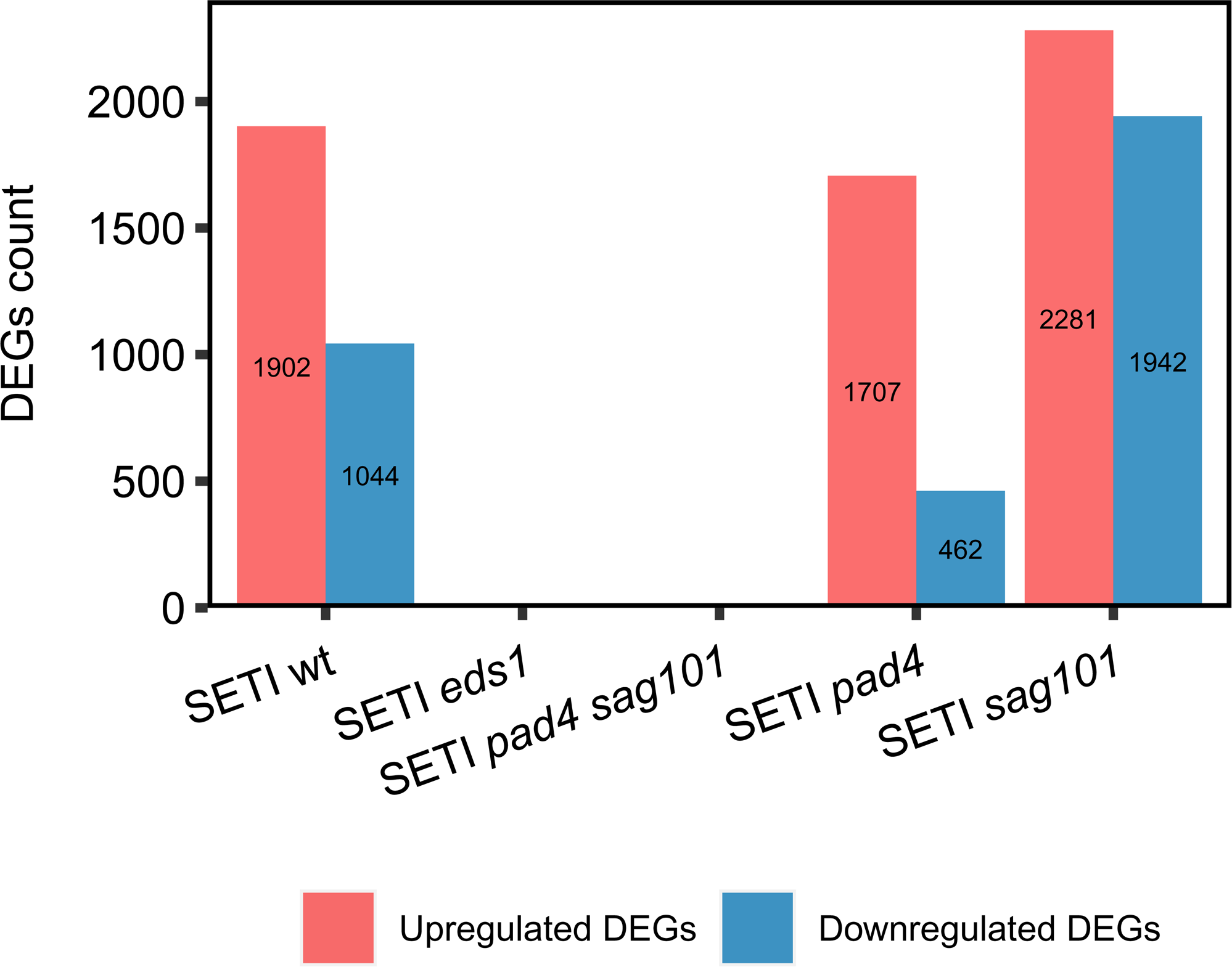
Number of up and down-regulated genes in all the plant lines after ETI-treatment induction. After ETI-treatment induction, the SETI_*sag101* line shows the highest DEG numbers compared to SETI_wt and SETI_*pad4*. However, SETI_*eds1* and SETI_*ps* didn’t show DEG after ETI induction. Hierarchical clustering was used to partition the DE genes into 10 clusters with 15 euclidean distance and ward.D clustering algorithm.

**Figure S5.**
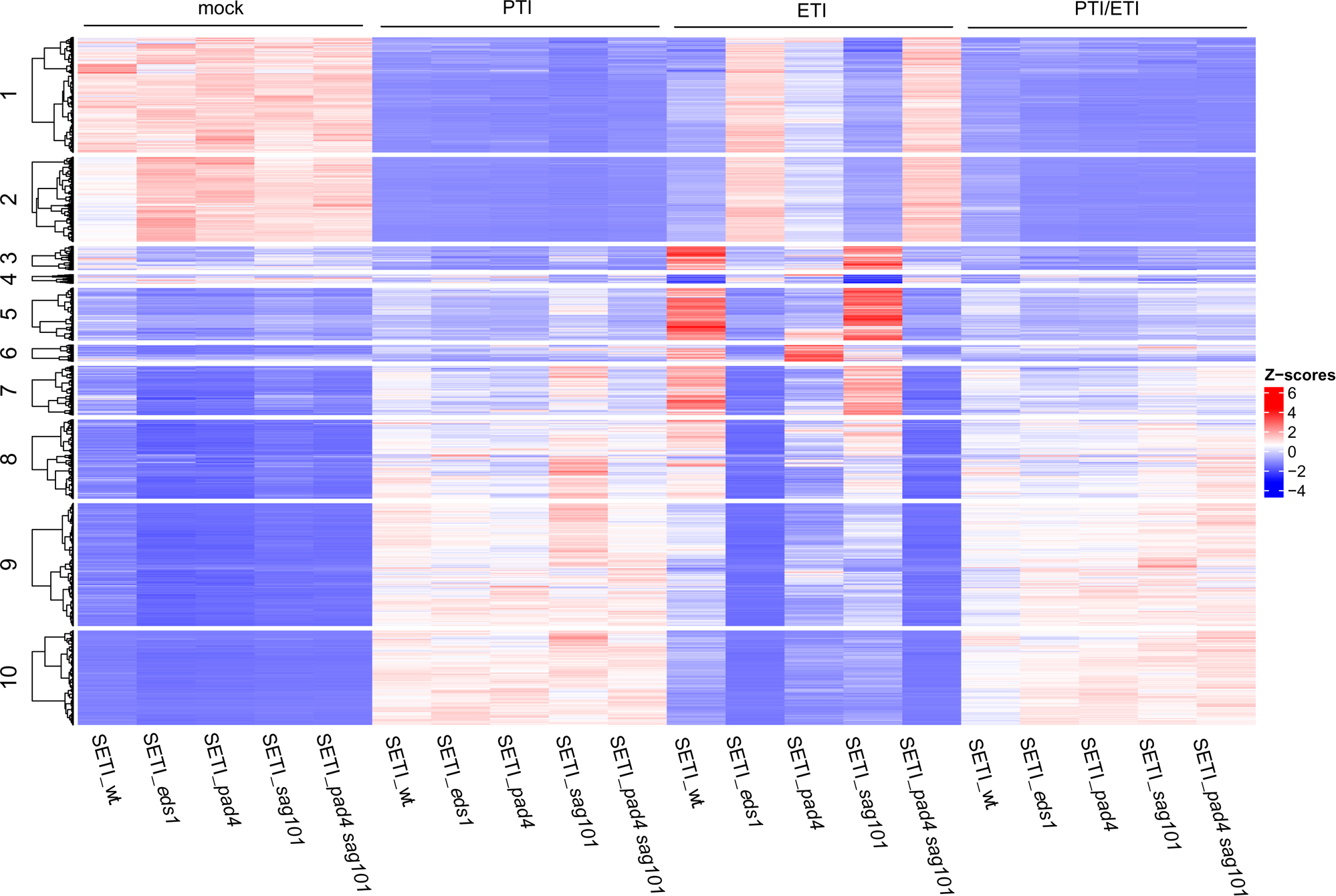
E2-induced expression changes differ between mutants. Five- to six-week-old SETI_wt, SETI_*eds1*, SETI_*pad4*, SETI_*sag101*, and SETI*_ps* plants were infiltrated with mock (10 mM MgCl**_2_**), PTI (*Pseudomonas fluorescens* (Pf0) in 10 mM MgCl**_2_**), ETI (50 uM E2 in 10 mM MgCl**_2_**), and PTI + ETI respectively.

**Figure S6.**
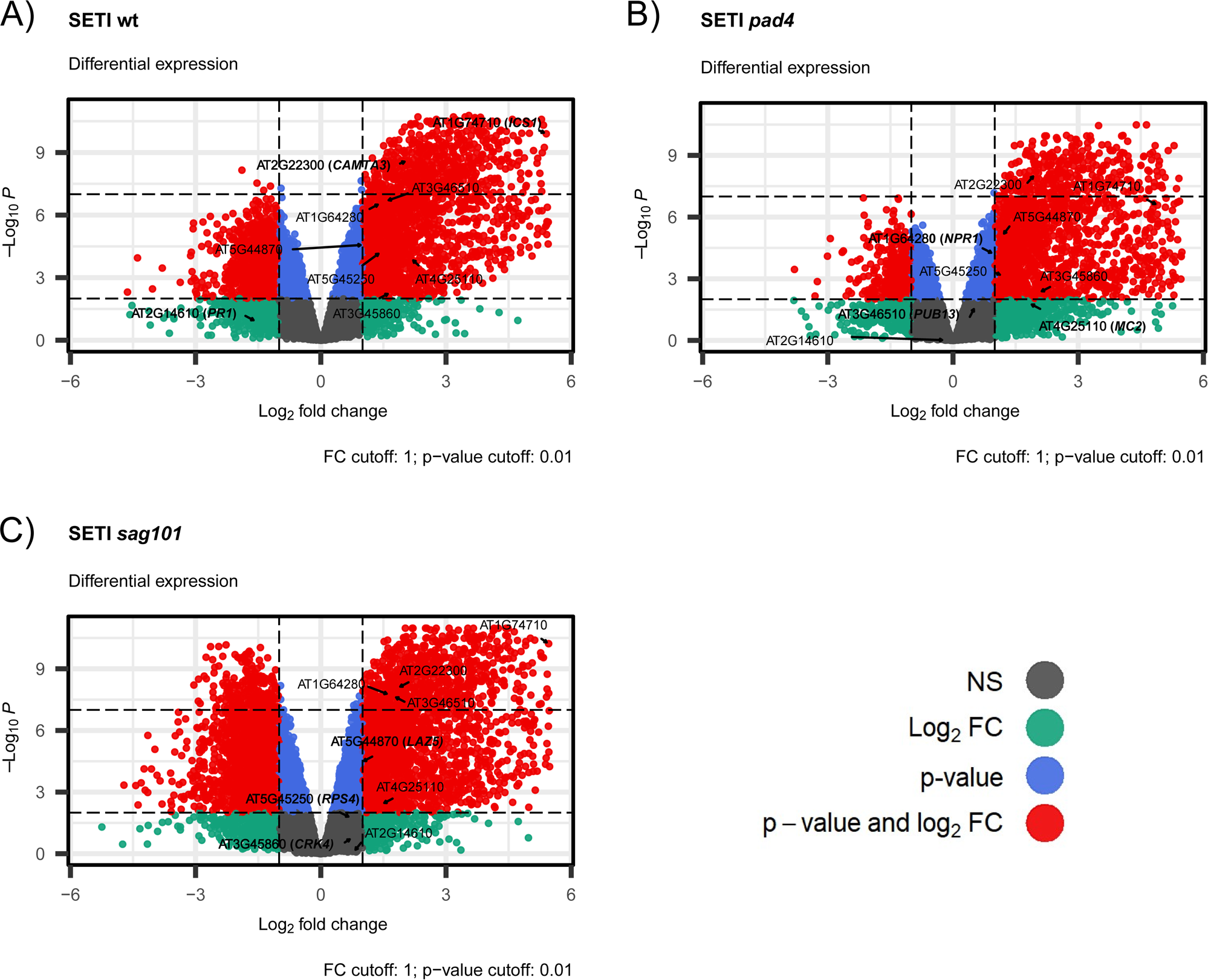
Volcano plot for SETI lipase-like protein mutants. Volcano plot showing expression changes between E2-vs mock-treated samples of SETI_wt plants. The gene names of three immune marker genes, three PAD4-dependent genes and three SAG101-dependent genes are highlighted here. Different colours highlighting different thresholds: red (p < 0.05, FC-cutoff < 1), blue (p < 0.05, FC-cutoff > 1), green (p > 0.05, FC-cutoff < 1), and grey (p > 0.05, FC-cutoff > 1).

**Figure S7.**
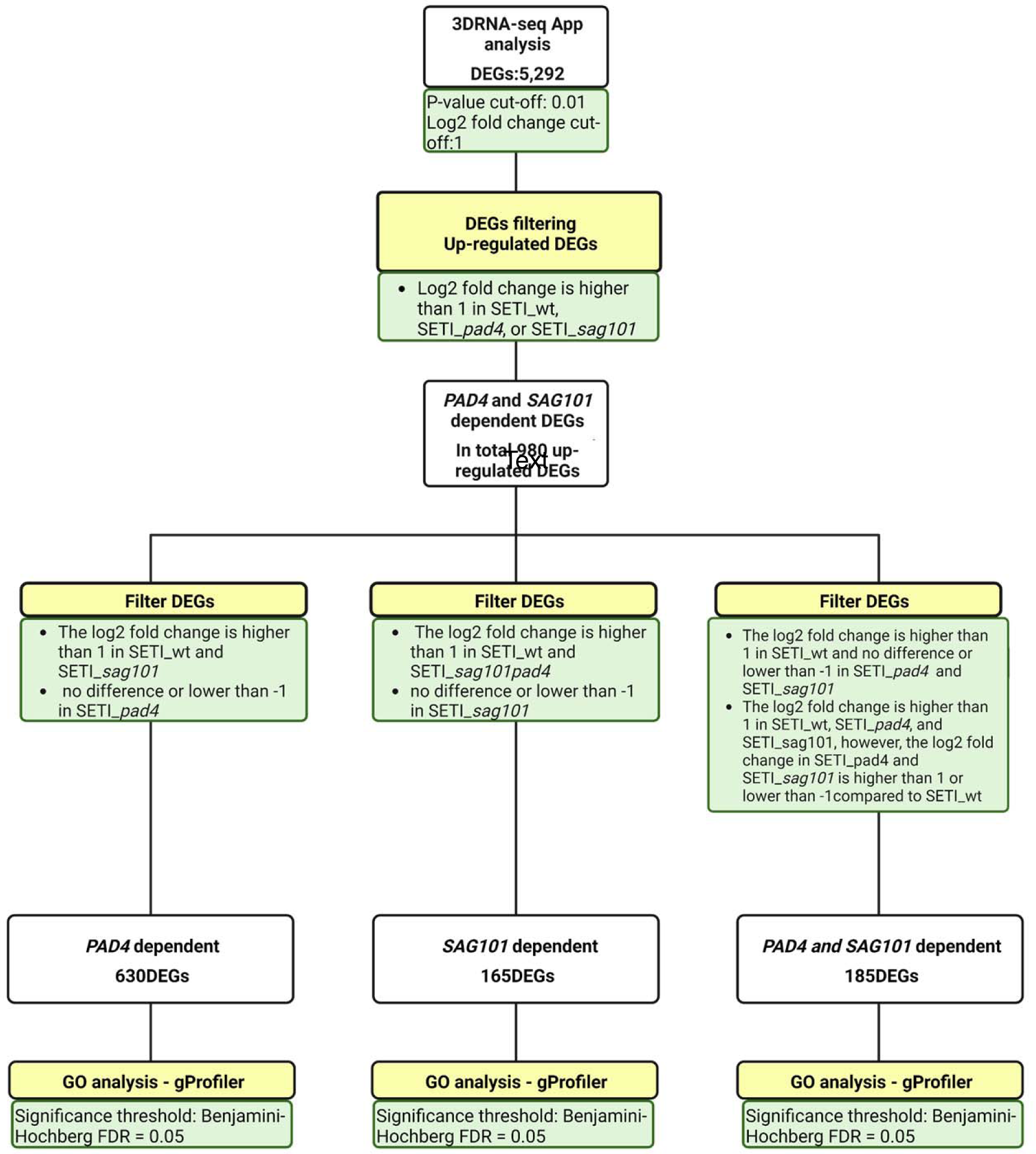
The workflow for RNA-seq analysis. We upload our raw data to the 3D RNA-seq app for rapid and accurate differential expression (DE), differential alternative splicing (DAS) gene and differential transcript usage (DTU) (3D) analysis^47,64^. The pipeline we chose is limma-voom; The p-value cut-off = 0.01 and the log2 fold change cut-off = 1. We got 5,292 DEGs and focused on up-regulated DEGs. We further classified these up-regulated DEGs into PAD4-dependent DEGs, SAG101-dependent DEGs, and PAD4/SAG101-shared DEGs. We processed GO term enrichment analysis by g:profiler with Benjamini Hochberg (BH) significance threshold FDR=0.05.

